# Measurement of activity of developmental signal transduction pathways to quantify stem cell pluripotency and phenotypically characterize differentiated cells

**DOI:** 10.1101/2021.04.14.439771

**Authors:** Yvonne Wesseling-Rozendaal, Laurent Holtzer, Wim Verhaegh, Anja van de Stolpe

## Abstract

Stem cell research is emerging both as a scientifically and clinically relevant area. One of the current challenges in stem cell research and regenerative medicine is assessment of the pluripotency state of induced pluripotent stem (iPS) cells. Once a stem cell differentiation process is initiated the challenge is how to assess the state of differentiation, and the purity of the differentiated cell population. Stem cell potency and differentiation states are determined by tightly coordinated activity of developmental signaling pathways, such as the Notch, Hedgehog, TGFβ, Wnt, PI3K, MAPK-AP1, and NFκB pathways. Source of the stem cells and culture protocols may influence stem cell phenotype, with potential consequences for pluripotency and in general for experimental reproducibility. Human pluripotent embryonic (hES) and iPS stem cell lines under different culture conditions, organ derived multipotent stem cells, and differentiated cell types, were phenotyped with respect to functional activity of developmental signaling pathways.

**Methods:** We previously reported on the development and validation of a novel assay platform for quantitative measurement of activity of multiple signal transduction pathways (STP) simultaneously in a single sample, based on interpreting a preselected set of target mRNA expression levels. Assays were used to calculate Notch, Hedgehog, TGFβ, Wnt, PI3K, MAPK-AP1, and NFκB signal transduction pathway activity scores for individual cell samples, using publicly available Affymetrix expression microarray data.

**Results:** Culture conditions (e.g. mouse versus human feeder) influenced pluripotent stem cell pathway activity profiles. hES and iPS stem cell lines cultured in the same lab under similar conditions showed minimal variation in pathway activity profile despite different genetic backgrounds, while across different labs larger variations were measured, even for the same stem cell line. Pathway activity scores for PI3K, MAPK, Hedgehog, Notch, TGFβ, and NFκB pathways rapidly decreased upon pluripotent stem cell differentiation, while increasing for the Wnt pathway. Further differentiation to intestinal progenitor cells resulted in higher PI3K, Wnt and Notch pathway activity. In multipotent intestinal crypt stem cells obtained from intestinal mucosa samples, similar Notch and even higher Wnt pathway activity were measured, which disappeared upon differentiation to mucosal cells.

**Conclusion:** Results support the validity of using these STP assays for quantitative phenotyping of stem cells and differentiated derivatives, and enabled definition of a pluripotency profile with high PI3K, MAPK, Hedgehog, TGFβ, Notch, and NFκB, and low Wnt pathway activity scores.

Measurement of combined signaling pathway activity scores is expected to improve experimental reproducibility and standardization of pluripotent and multipotent stem cell culture and differentiation. It enables controlled manipulation of signaling pathway activity using pathway targeting compounds. An envisioned additional utility may lie in quality control for regenerative medicine purposes.

## Introduction

Generation of human embryonic stem (hES) cell lines preceded the emergence of induced pluripotent stem (iPS) cell technology [1]. Nowadays, well described and established protocols enable the generation of a pluripotent stem cell line from a variety of cell types of a healthy or diseased individual [2]. Clinical use for regenerative medicine purposes, as well as creation of in vitro healthy and diseased tissue models to improve understanding of disease pathophysiology and drug development, are major stem cell applications. Especially for iPS cells, the generation of cell types for tissue repair holds great clinical potential for treating and potentially curing a multifold of diseases [1],[4],[5],[6],[7]. Pluripotent stem cell culture and differentiation towards specific cell types is of increasing importance for the creation of in vitro disease models. Since iPS-based disease models are generated based on the genetics of the donor and therefore unique for each patient, utility may also lie in a personalized disease model for patients with certain (lifelong) inherited diseases [7].

All these applications face significant challenges. For example, for in vitro disease models it is important that they are sufficiently reproducible and representative for in vivo human disease. For regenerative medicine, a major challenge is to reproducibly generate the required differentiated cells with sufficient purity [8].

Unfortunately, despite similar cellular morphology, pluripotency - defined as the capability to differentiate into the three germ layers: ectoderm, endoderm, and mesoderm - may differ between various hES and iPS cell lines [9]. Observed differences in pluripotency have yet to be related to cellular mechanisms that determine differentiation potential and can be adjusted in cell culture. In addition, assessment of cell differentiation state and population purity mostly relies on the introduction of reporter genes, specific for a required differentiation or maturation state [10],[11],[12].

Consequently, there is a need for methods to better characterize the different types of stem cells and derivatives with respect to pluripotency and differentiation state, preferably in a quantitative manner [13],[14],[15]. Both pluripotency and stem cell differentiation are controlled by coordinated activity of a number of highly conserved signal transduction pathways [3],[17],[18],[19],[20]. They can be roughly categorized as hormonal driven nuclear receptor pathways (e.g. androgen and estrogen receptor pathways), core developmental pathways (e.g. Wnt, Hedgehog (HH), TGFβ, and Notch pathways), the NFκB pathway, and the growth factor regulated signaling pathway network, that is, the PI3K-AKT-mTOR, MAPK and JAK-STAT pathways [18]. In addition to embryonic development, they regulate many physiological processes, while in diseases control of signaling pathway activity is typically altered or lost [21]–[33].

Assessment of combined activity of these signaling pathways was expected to provide relevant information on (pluri)potency or differentiation state of cells. We previously described a novel assay platform to quantitatively measure the functional activity of a number of signal transduction pathways (STP) in individual cell and tissue samples, based on the inference of pathway activity from measurements of mRNA levels of target genes of the transcription factor associated with the signaling pathway [16],[36],[37],, [36]. To define a pluripotency-associated pathway activity profile, and to address the potential of the STP assays to increase experimental reproducibility, we determined quantitative signal transduction pathway activity profiles of different types of hES and iPS cell lines and differentiated derivatives across different labs, as well as primary organ-derived stem cells.

## Methods

### Development and biological validation of MAPK-AP1 pathway activity assay

The method to develop signal transduction pathway assays has been described before [16],[37]. Here we report development and biological validation of an additional assay for measuring the activity of the MAPK-AP1 signaling pathway. In brief, a Bayesian computational network model for a signal transduction pathway is created to infer the probability that the pathway-associated transcription factor is in a transcriptionally active state. The Bayesian network describes (i) the causal relation that a target gene is up- or downregulated depending on the transcription complex being active or inactive and (ii) the causal relation that a signal from a gene expression microarray (e.g. Affymetrix Human Genome U133 Plus 2.0) probeset is high or low depending on the target gene being up or down. The list with target genes used to develop the new MAPK-AP1 pathway assay is available in the Supplementary Materials and Methods (Supplemental Table S1).

### Analysis of signal transduction pathway activity on Affymetrix expression microarray data

Affymetrix Human Genome U133 Plus 2.0 microarray data of stem cell studies were downloaded from the public GEO database (https://www.ncbi.nlm.nih.gov/gds) and signaling pathway activity scores for AR, ER, PI3K-FOXO, MAPK-AP1, HH, Notch, TGFβ, Wnt, and NFκB pathways were calculated as described before [37]. For each signaling pathway, pathway activity is inferred from the log2 odds that the pathway is active. For the PI3K pathway, activity of the FOXO transcription factor is calculated, and PI3K pathway activity can be inferred from the FOXO transcription factor activity score by inverting it, on the premise that no cellular oxidative stress is present, as described before [35], [38]. Quality assessment of Affymetrix data was performed prior to pathway analysis as described [37]. Shown are results of respective pathway activity scores in a multi-pathway analysis of data from individual samples.

### Interpretation of signal transduction pathway activity scores [37]

An important and unique advantage of the pathway activity assays is that they can in principle be performed on each cell type. Important considerations for interpretation of log2 odds pathway activity scores are:

1. on the same sample log2 odds pathway activity scores cannot be directly compared between different signaling pathways, since each of the signaling pathways has its own range in log2 odds activity scores;
2. the log2 odds range for pathway activity (minimum-maximum activity) may vary depending on cell type. Once the range has been defined using samples with known pathway activity, on every new sample the absolute value can be directly interpreted against that reference. If the range has not been defined, only differences in log2 odds activity score between samples can be interpreted;
3. pathway activity scores are highly quantitative, and even small differences in log2 odds can be reproducible and meaningful;
4. a negative log2 odds ratio does not necessarily mean that the pathway is inactive.

### Analyzed public GEO datasets containing Affymetrix HG-U133 Plus 2.0 expression microarray data

#### Pluripotent stem cells

- Dataset GSE19902 contains data from human embryonic stem cells (hES-T3), cultured using different culture protocols (with 4 ng/ml bFGF): on autogeneic fibroblast feeder layer derived from hES-T3 cells or its conditioned medium; on murine embryonic fibroblast (MEF) feeder layer or its conditioned medium [39].
- Dataset GSE52658 contains data from hES cells, adapted to feeder-free culture on Matrigel-coated plates with MEF-conditioned medium with 15 ng/ml FGF2 [40].
- Dataset GSE7879 contains data from mesenchymal stem cell-derived cells (VUB01; hES), cultured on MEF feeder, with 4 ng/ml bFGF, and SA01 (Cellartis), or cultured on human foreskin fibroblast feeder, supplemented with 4 ng/ml bFGF [41].
- Dataset GSE9440 contains data from hES cells cultured on MEF, supplemented with 4 ng/ml bFGF [42].
- Dataset GSE17312 contains data from iPS and hES cell lines, no information on culture conditions reported (*BI Human Reference Epigenome Mapping Project*) [43], [44].
- Dataset GSE74358 contains data from four different iPS cell lines cultured on murine embryonic fibroblast (MEF) feeder layer (with 10 ng/ml bFGF) [45].

#### Organ-derived multipotent stem cells

- Dataset GSE84500 contains data from human bone marrow mesenchymal stem cells [46].
- Dataset GSE31255 contains data from colon epithelial cells microdissected from healthy colon tissue biopsies, including intestinal crypt stem cells [47].

#### Differentiated stem cell derivatives

- Dataset GSE52658 contains data from the differentiation of hES cells to mesoderm and endoderm [40].
- Dataset GSE74358 contains data from iPS cell differentiation to ectodermal neuronal progenitors and neuronal cells, 4 different iPS cell lines [45].

### Statistics

Two-sided Wilcoxon signed-rank statistical tests were performed, as indicated in the respective figures. In view of the small sample sets, extensive statistical analysis was considered as not useful. Also, with very low sample numbers (n<3) visualization and descriptional analysis of results (based on non-overlapping scores +/-SD) is performed, since statistical analysis is not informative.

## Results

### Development of a MAPK-AP1 pathway activity assay

Ligand-activated Receptor Tyrosine Kinase Receptors, integrin receptors, and G-protein-coupled Receptors (GPCRs) can activate the MAPK pathway, consisting of sequential activation of a number of kinases (MAP-Kinase-Kinase-Kinase, MAP-Kinase-Kinase, and a MAP-Kinase), resulting in activation of the AP1 transcription factor consisting of dimers between members of the Fos and Jun protein family [48]. The target genes used in the MAPK-AP1 pathway model are listed in Supplementary table S1. In in vitro culture experiments, 12-O-Tetradecanoylphorbol-13-acetate (TPA; also referred to as phorbol-12-myristate 13-acetate, PMA) is the archetypal activator of AP1, and both calibration and validation samples for the pathway model were based on the use of this activator molecule in various cell types. For calibration purposes vehicle-treated (MAPK-AP1 inactive) and 500 nM TPA-treated (MAPK-AP1 active) HepG2 liver cell samples [49] were used (Supplementary Figure S1A).

After freezing the model, independent samples were used for validation of the model (Supplementary Figure S1B-E). The MAPK-AP1 model correctly measured increased AP1 activity in 100 nM TPA-treated U937 cells [50] (Supplementary Figure S1B), while knockdown of ZXDC1, a transcription factor regulating gene transcription during myeloid cell differentiation after AP1 activation, did not change MAPK-AP1 activity in TPA-treated U937 cells, confirming that the target genes in our MAPK-AP1 model are specifically activated by MAPK-AP1. In other experiments, U937 cells had been treated with 50 ng/ml TPA [51], and K562 (erythroleukemia) cells with 10 nM and 100 nM TPA/PMA respectively [52], [53], resulting in increased MAPK-AP1 pathway activity scores (Supplementary Figure S1C and S1D). The MAPK-AP1 model was further validated on samples treated with the protein kinase C activating drug PEP008 (20-O-acetyl-ingenol-3-angelate), and with effects comparable to TPA. SK-MEL-5 (melanoma; Supplementary Figure S1E) cells and COLO-2005 (colon cancer; Supplementary Figure S1F) cells were treated with 1 μg/mL PEP008 or 1000 ng/mL TPA (only the SK-MEL-5 cells) [54], resulting in increased MAPK-AP1 pathway activity.

Finally, the MAPK-AP1 pathway model was evaluated on tamoxifen-resistant MCF7 human breast cancer cells treated 10 nM growth hormone heregulin (Supplementary Figure S1G) to activate the MAPK-AP1 signaling pathway [55]. Oyama et al. reported that MAPK activity is altered in tamoxifen-resistant MCF7, which is reflected by the strong increase in MAPK-AP1 pathway activity upon heregulin stimulation compared to wild-type MCF7.

### Measuring signal transduction pathway activity in various stem cell lines and differentiated derivatives

Signaling pathway analysis was performed on Affymetrix expression microarray data from various types of stem cells, cultured in different labs, and either pluripotent, multipotent or differentiated in a specific direction. In general, between samples from the same pluripotent stem cell line or between different cell lines cultured in the same laboratory, pathway activity scores showed relatively small variations, but across laboratories larger differences in pathway activity profiles were observed. Pathway activity profiles clearly differed between pluripotent, multipotent and differentiated cells, and were influenced by culture conditions. Aside from providing insights into the phenotype of the investigated stem cell types, the presented results also emphasize the quantitative nature of the pathway assays (Figure 1-4).

**Figure 1.**
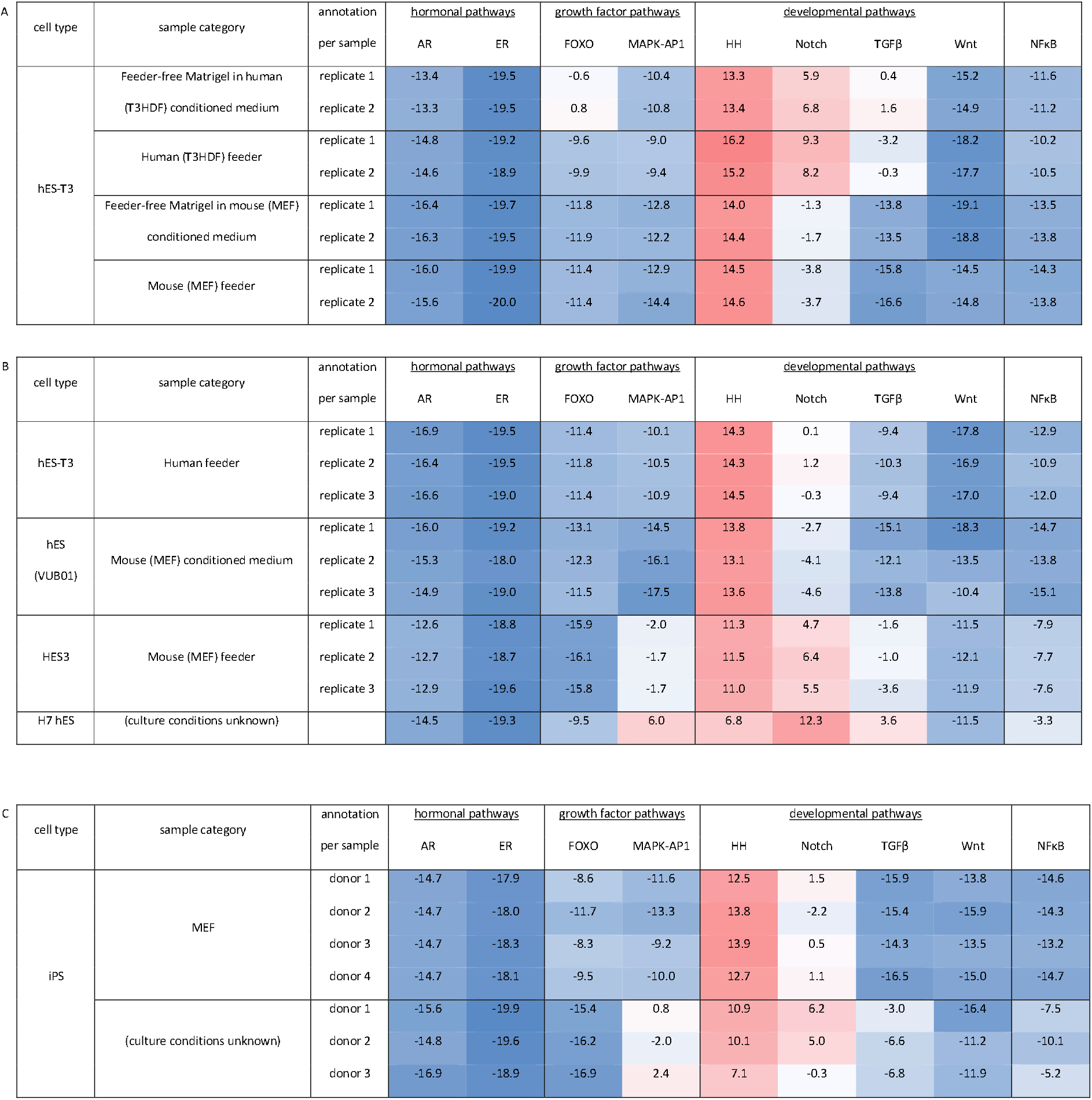
Human stem cell characterization in terms of signaling pathway activity. Shown are results of respectively androgen receptor (AR), estrogen receptor (ER), PI3K-FOXO, MAPK-AP1, Hedgehog (HH), Notch, TGFβ, Wnt, and NFκB pathway activities for individual samples; pathway activity is color coded ranging from blue (most inactive) to red (most active) [84]. The activity of the FOXO transcription factor is measured as the inverse readout for the activity of the PI3K pathway, in the absence of oxidative stress [35]. Individual sample annotation as provided in the public GEO database is on the left. A. hES-T3 cells. GEO dataset GSE19902 containing sample data from pluripotent human embryonic stem cells (hES-T3), cultured under different conditions, either on a human feeder layer (T3HDF) or with T3HDF-conditioned medium (supplemented with 4 ng/ml bFGF), or on a mouse feeder layer (MEF) or with MEF-conditioned medium (containing 4 ng/ml bFGF) [39]. B. hES cells. GEO dataset GSE84500, containing sample data from datasets GSE9440 and GSE35603. Undifferentiated hES-T3 (Cellartis) cultured on human foreskin fibroblasts supplemented with 4 ng/ml bFGF, and hES (VUB01) cells cultured on MEF [42],[46]. GSE52658, HES3 cells cultured with MEF-conditioned medium with 15 ng/ml FGF2 [40]. GSE17312, H7 hES cell line, from BI Human Reference Epigenome Mapping Project (culture conditions unknown) [43]. C. Human iPS cells. GSE74358, iPS cell lines cultured on MEF (with 10 ng/ml bFGF). iPS lines obtained from Amish pedigree [45]; GSE17312, iPS cell lines from BI Human Reference Epigenome Mapping Project (culture conditions unknown) [43].

#### STP activity scores in hES cells vary with different culture conditions

Signaling pathway activities were compared between hES-T3 human embryonic stem cells cultured in different ways: (1) in conditioned medium of the human feeder layer, supplemented with 4 ng/ml bFGF; (2) on a human autogeneic fibroblast feeder layer; (3) in MEF-conditioned medium, supplemented with 4 ng/ml bFGF; and (4) on a conventional murine MEF feeder layer. HH pathway activity scores were indicative of an active pathway and relatively independent of culture conditions (Figure 1A). Pathway activity scores were consistently very low for the ER, AR, Wnt, and NFκB pathways. When comparing human with murine culture conditions (either on feeder layer or with conditioned medium), higher pathway activity scores of the Notch and TGFβ pathways and to a lesser extent of the MAPK-AP1, and NFκB pathways, and lower PI3K (the inverse of higher FOXO transcription factor pathway activity – see Methods) pathway activity was measured in cells cultured in a human environment. The impact of switching from culturing on a feeder layer to culture in conditioned medium on signaling pathway activity was small compared to the different effects of human and mouse culture conditions. Cells cultured on a human feeder had higher PI3K and Notch pathway activity scores compared to a feeder-free human conditioned medium.

#### STP activity scores in pluripotent hES cell lines differ between laboratories

Data from three independent studies were available in which undifferentiated hES cell lines had been cultured on MEF or in MEF-conditioned medium (Figure 1B). For the H7 hES cell line, analyzed within the *BI Human Reference Epigenome Mapping Project*, culture conditions were unknown. For all cell lines, pathway activity profiles were highly similar between replicate cultures in the same lab/study, but varied depending on the lab where cells had been cultured. The H7 hES cell line (of which culture conditions were unknown) had a clearly different pathway activity profile, with the highest MAPK-AP1, Notch, TGFβ, NFκB pathway activity scores, the relatively low HH and PI3K (higher FOXO activity) pathway activity score.

For the same hES-T3 cell line, data were available from two independent studies from different labs (Figure 1A and B), using two different human feeders. Notch and TGFβ pathway activity scores were much lower in cells cultured on human foreskin fibroblasts (figure 1B), compared to culture on an autogeneic human feeder (Figure 1A). These results provide more evidence that even seemingly small differences in culture protocol may affect the hES cell phenotype, and may be responsible for differences in experimental results between laboratories.

#### STP activity scores are similar for different iPS cell lines under the same culture conditions, but differ between labs

STP activity profiles for iPS cell lines obtained from four different donors and cultured in the same lab, under the same culture conditions, showed relatively minor differences (Figure 1C, dataset GSE74385). However, relatively large differences were found between these iPS cell lines and the three iPS cell lines in the second dataset (GSE17312) for which culture protocol(s) were unknown. Notably, the GSE74385 dataset was from the Old Order Amish group with high incidence of bipolar disorder, and genetic backgrounds may have been quite similar. Thus, observed differences in STP activity profiles between the two sets of iPS cell lines may have been caused by substantial genetic differences, or by differences in iPS cell line generation protocol, or by different culture protocols.

#### Phenotypic comparison between different stem cell lines

No consistent differences in STP activity profiles between the various hES cell lines, or between hES and iPS cell lines, were found (Figure 1). However, genetically determined differences, or differences originating from different protocols used to establish the cell line, may have been overshadowed by major effects of culture conditions on the phenotype.

### Pluripotent stem cell differentiation to endodermal, mesodermal, and ectodermal derivatives

#### hES cell differentiation to endodermal and mesodermal derivatives

Pluripotent stem cells are by definition able to differentiate to the three germ layer lineages. To reliably assess the pluripotency of stem cells, it is important to define the changes in signaling pathway activity associated with early differentiation and loss of pluripotency. Loh et al. differentiated hES-3 cells in defined medium into mesoderm and definitive endoderm, and subsequently to anterior foregut as the precursor for lung and thyroid, to posterior foregut as the precursor for pancreas and liver, and to mid/hindgut as the precursor for intestinal tissue [40]. Signaling pathway activity scores were measured during the different stages in the differentiation process (Figure 2). Pluripotent hES3 cells (also shown in Figure 1B, GSE52658) showed a pathway profile with relatively high MAPK-AP1, PI3K (i.e. low FOXO activity), HH, TGFβ and Notch pathway activity scores, while Wnt pathway activity scores were typically low. Initial differentiation to anterior primitive streak (APS, day 1, precursor for mesoderm and definitive endoderm) resulted in a reduction in activity of the pathways already associated with pluripotency in the datasets described above, that is, the HH, Notch, TGFβ, NFκB, MAPK-AP1 and PI3K (increase in FOXO activity) pathways, with a simultaneous increase in Wnt pathway activity. Differentiation from APS to definitive endoderm (DE, day 3) caused a decrease in Wnt, a further decrease in PI3K (i.e. increase in FOXO activity) pathway activity, and a simultaneous increase in MAPK-AP1 and NFκB pathway activity. When instead the APS cells were differentiated to mesoderm (day 3), cells differed from DE by a higher Wnt and TGFβ pathway activity and lower NFκB pathway activity scores. Subsequent differentiation to mid/hindgut as intestinal precursor cells, showed a distinct STP activity profile with high PI3K, Wnt, Notch, and low TGFβ pathway activity scores. Clearly, already after one day of differentiation, some pluripotency-associated signaling pathways have lost activity – that is, the MAPK-AP1, PI3K, HH, Notch, TGFβ, and NFκB pathways – while the Wnt pathway increased in activity.

**Figure 2.**
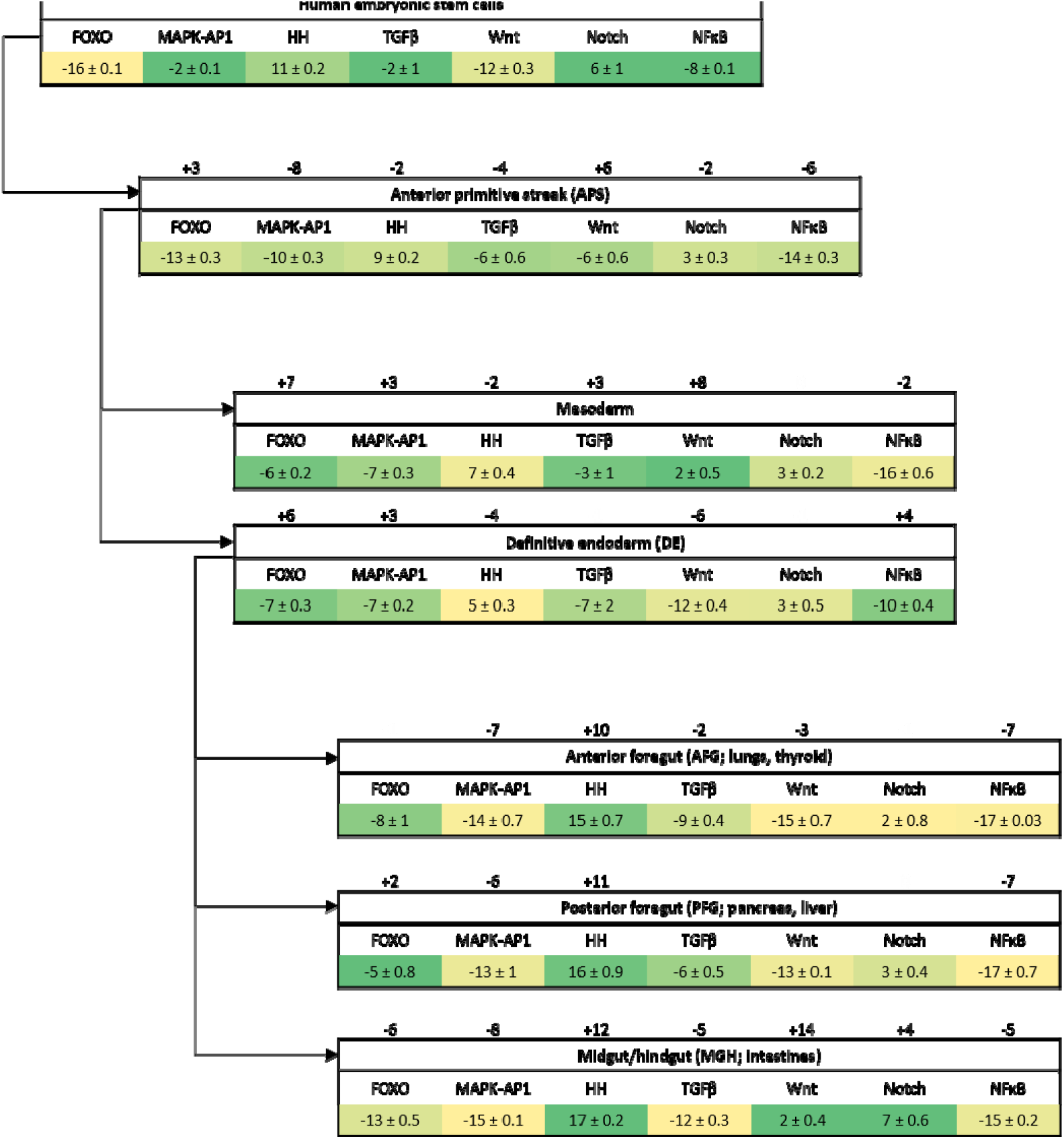
Characterization of cells in various stages of differentiation in terms of the activity of multiple signaling pathways (presented as mean ± standard deviation, n=3 replicates). Using public mRNA expression datasets (GSE52658), we have measured combined pathway activity on the stem cells during different stages in the differentiation process (as listed in detail in Supplementary Figure S2). Color-coding is used to visualize changes in signaling pathway activity scores during differentiation: ranging from dark green (indicating the highest observed activity in this set for each pathway separately) to yellow (indicating the lowest observed activity in this set for each pathway separately). Loh et al. differentiated human embryonic stem cells (HES) into definitive endoderm and mesoderm, and subsequently to anterior foregut as the precursor for lung and thyroid, to posterior foregut as the precursor for pancreas and liver, and to mid/hindgut as the precursor for intestinal tissue [40]. hES cells were cultured under serum-free conditions in defined medium. To obtain anterior primitive streak cells (APS), ectoderm differentiation was excluded by activation of TGFβ and Wnt pathway activity, while inhibiting the PI3K/mTOR pathway. The differentiation step from APS to definitive endoderm (DE) consisted of stimulation with high Activin and BMP blockade to prevent the formation of mesoderm. Subsequently, DE cells were differentiated for four days to different types of foregut cells (AFG/PFG/MHG), using treatment with BMP, Wnt, and FGF.

#### Differentiation of iPS cells to ectoderm-derived neuronal cells

To investigate changes in signaling pathway activity during differentiation to the ectodermal direction, a dataset was available in which iPS cells from different donors had been differentiated towards neuronal cells. Initial (two weeks) differentiation to neural progenitor cells was associated with reduced MAPK-AP1 pathway activity scores, while Wnt pathway activity increased (Figure 3 and Supplementary Figure S3). Subsequent differentiation to early and late stage neurons was associated with a reduction in PI3K (increased FOXO activity scores) and HH pathway activity, and increased Notch pathway activity.

**Figure 3.**
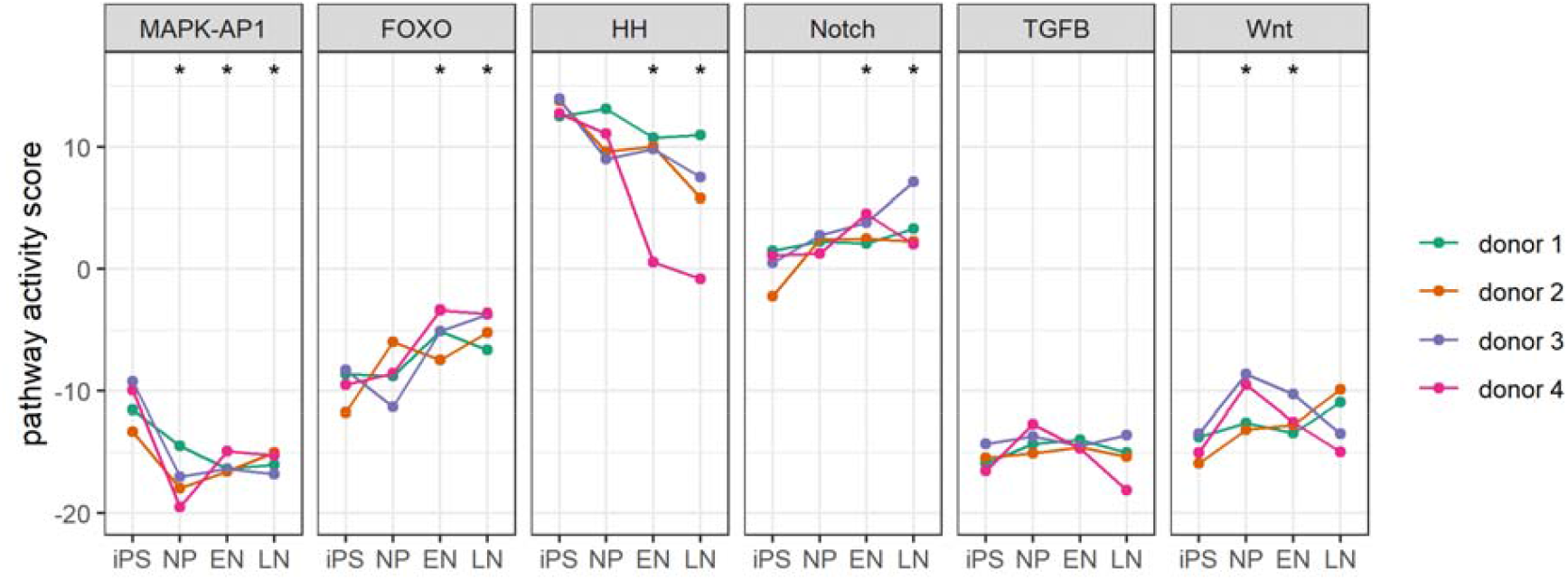
iPS cell differentiation to neuronal progenitor (NP) cells and early (EN) and late neurons (LN). iPSCs, NPs and neurons matured in culture for 2 weeks (EN) or 4 weeks (LN). iPS cell lines were derived from the Old Order Amish group. Sample data from dataset GSE74358 [45]. Statistically significant differences (Wilcoxon two-sided test) – compared to the iPS group – are indicated with an asterisk (p<0.05).

### Signaling pathway analysis of organ-derived (bone marrow and intestinal mucosa) multipotent stem cells

Primary organ-derived stem cells are not pluripotent but at best multipotent [1]. Two types of organ-derived stem cells were analyzed: from bone marrow (mesodermal) and from intestinal crypts (endodermal) (Figure 4).

**Figure 4.**
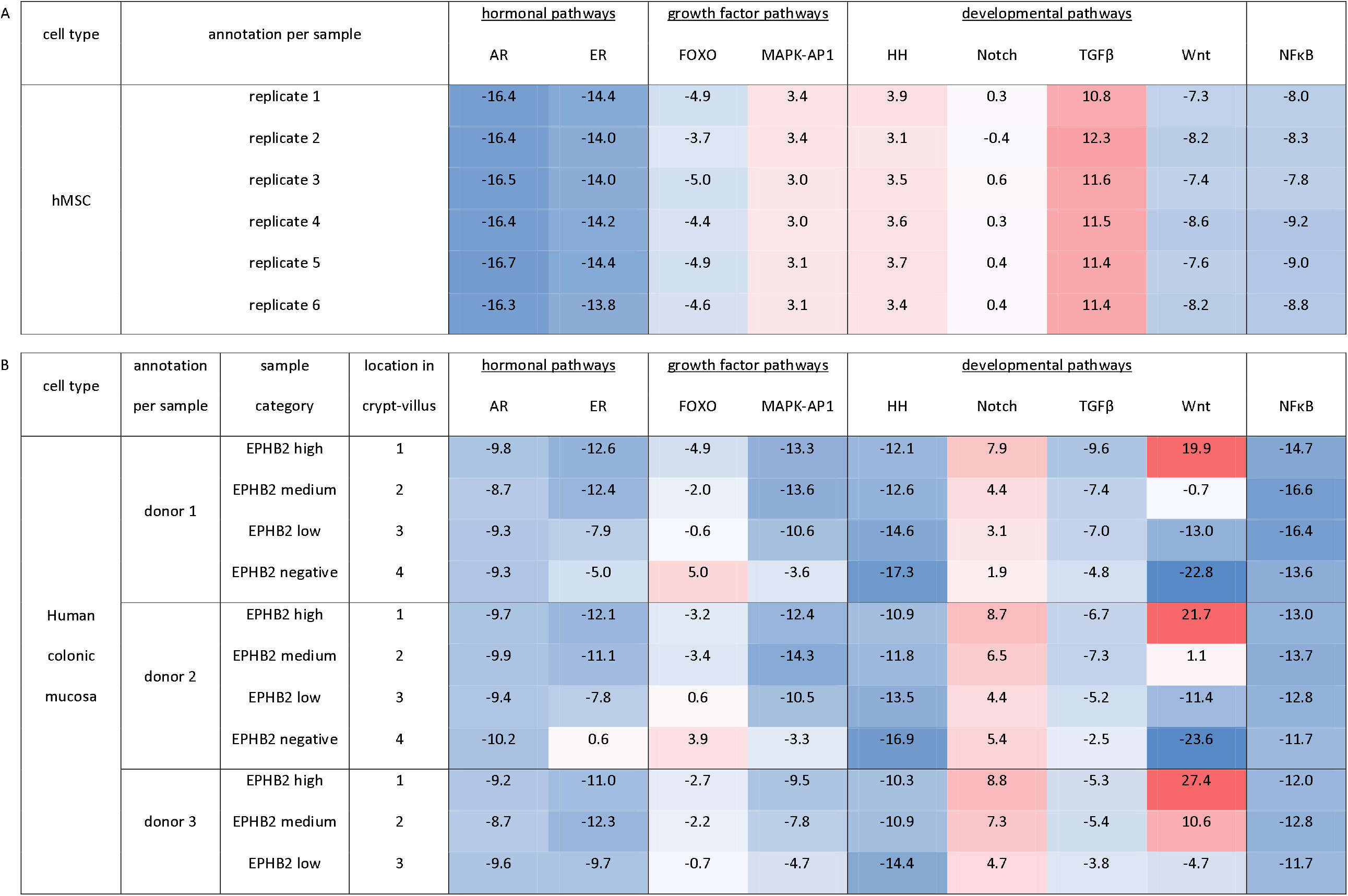
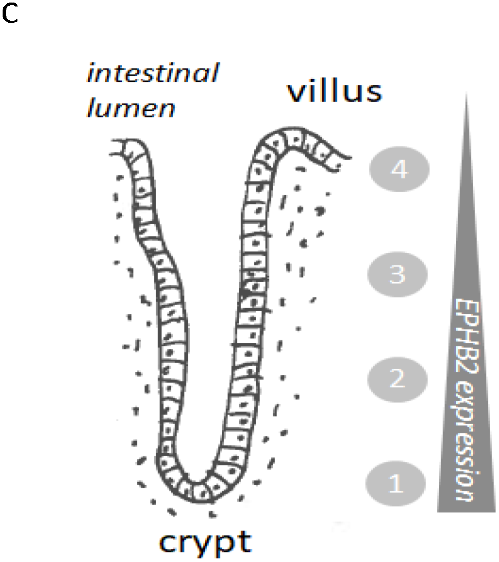
Organ-derived stem cells A. Human bone marrow-derived mesenchymal stem cells, sample data from GEO dataset GSE84500 [46]. B. Normal human colonic mucosa cells from three different crypt-villus locations were isolated from colon biopsies from three individuals (dataset GSE31255). Highest ephrin type-B receptor 2 (EPHB2) protein levels characterize intestinal stem cells. Sampling locations are indicated (roughly): 1, high EPHB2; 2, medium EPHB2; 3, low EPHB2; 4, negative EPHB2 [47]. C. Intestinal crypt-villus structure, with decreasing EPHB2 expression from crypt compartment containing intestinal stem cells, to villus with differentiated mucosa cells. Sampling locations are indicated as 1-4.

Activity of the HH pathway was lower in both these stem cell types comparable to the activity seen in pluripotent hES and iPS cell lines. In addition, cultured human mesenchymal stem cells from bone marrow had a higher MAPK-AP1 and TGFβ pathway activity compared to iPS and hES stem cells (Figure 4A).

Intestinal crypts contain a small number of stem cells, which differentiate while migrating towards the villus along a decreasing EPHB2 gradient [52],[61]. For the analyzed study, colon mucosa samples had been obtained from four locations along the crypt-villus axis. Crypt stem cells (indicated with 1 in Figure 4B, 4C) showed a combination of high Wnt, Notch, and PI3K pathway activity, with pathway activity scores decreasing upon differentiation (indicated with 2,3,4) towards the villus mucosa, while MAPK-AP1 and ER pathway activity increased towards the villus (Figure 4B). Among the various types of analyzed stem cells, only the crypt stem cells exhibited a high Wnt and to a lesser extent Notch, pathway activity, which was rapidly lost upon differentiation. The pathway activity profile seen in the intestinal mucosa stem cells (crypt cells) closely resembles that observed in the mid/hindgut cells (Figure 2), which are considered progenitors for intestinal mucosa cells, although in the crypt stem cells Wnt pathway activity is even higher.

### Relation between STP activity scores and functional activity of signal transduction pathways

As described in the Method, pathway activity scores can only be interpreted towards an active or inactive pathway in an absolute manner when the range in pathway activity scores (between minimal and maximal functional pathway activity) in the cell type is known and therefore can serve as a reference. The here presented STP analysis helps to establish such reference levels, based on the results of cells with a known pathway activity.

The Wnt pathway should be active in intestinal crypt stem cells, but inactive in pluripotent stem cells [56],[57]. In assumed pluripotent stem cells Wnt pathway activity scores were between -19 and -10, representing an inactive Wnt pathway (Figures 1-2). In multipotent intestinal crypt stem cells with a (highly) active Wnt pathway, Wnt pathway activity scores were between +20 and +27; towards the villus the pathway becomes increasingly inactive, resulting in Wnt pathway activity scores between -24 and -5. Using these results, while realizing the limitation of small sample numbers, we can now set a putative range for Wnt pathway activity scores in stem cells and derivatives between around -24 (inactive) and +28 (highly active). Subsequently we can use this reference range and conclude that the Wnt pathway was probably moderately active in midgut/hindgut cells (Figure 2). For the other pathways, the ranges in pathway activity scores could also be established based on analysis of the various undifferentiated and differentiated cell types (PI3K-FOXO: between -17 and +5; MAPK-AP1: between -20 and +6; HH: between -17 and +17; Notch: between -5 and +12; TGFβ: between -18 and +12; NFκB: between -17 and -3) (Figures 1-4, S2, S3). These ranges are much larger than the variations measured in the same cell types under the same culture conditions, or even between different iPS cell lines. This indicates that changes in functional signaling pathway activity occur with changing culture protocols and differentiation of stem cells. However, in the absence of sample sets with known pathway activity the range only represents a minimum range, and only relative differences in pathway activity can be inferred.

## Discussion

In order to define a signaling pathway activity profile for stem cell pluripotency, in pluripotent hES and iPS cells and differentiated derivatives, as well as in organ-derived multipotent stem cells, activity was measured of signal transduction pathways known be important for self-renewal and differentiation. A previously described, validated set of signal transduction pathway assays was used to calculate pathway activity scores for the AR, ER, PI3K, Wnt, TGFβ, HH, Notch, and NFκB pathways using RNA expression data [16],[35],[37],[36]. To enable a more complete analysis of signaling pathway activity, for the MAPK-AP1 pathway an additional assay was developed, biologically validated on a number of different cell types, and added to the existing signaling pathway assay platform.

### Influence of stem cell culture protocol on pluripotency phenotype

Comparing signaling pathway activity scores between pluripotent hES cells and iPS cell lines across different labs revealed that STP activity profiles were highly reproducible, even between different stem cell lines, within a lab and when cultured with the same protocol, but showed relatively large variations across laboratories. This was nicely illustrated by the hES-T3 cell line: a single cell line, cultured in two different labs under seemingly quite similar conditions, but resulting in very different Notch and TGFβ pathway activity scores, possibly determined by the type of human feeder that was used. On the other hand, iPS cell lines from different donors with different genetic backgrounds, cultured in the same lab with the same protocol, showed very comparable pathway activity scores. These results emphasize the important influence of culture protocols on pluripotent stem cell phenotype.

While determination of the influence of specific culture conditions on pathway activity was hindered by many unknowns in culture protocols across different labs, one hES dataset from a single lab provided some initial insight. Here, the pluripotency phenotype in terms of pathway activity appeared to be especially influenced by a murine versus human environment, where a human culture environment was associated with higher pathway activity scores for especially the Notch, TGFβ, and MAPK-AP1, pathways, and lower PI3K pathway activity scores. An explanation for signaling pathways being more active in a human culture environment may be availability of human ligands for the specific pathway receptors instead of murine ligands, where human ligands may more effectively activate the signaling pathway. However, aside from species differences, many other culture protocol-associated factors may influence pathway activity profiles across labs, for example variations in the constituents of defined culture medium. Signaling pathway activity can be induced dose-dependently by a specific ligand present in culture medium, and can subsequently influence activity of other signaling pathways through crosstalk. For example, the MAPK-AP1 pathway can be activated by basic Fibroblast Growth Factor (bFGF), but activity of this pathway can also be modified by crosstalk with other signaling pathways, notably the TGFβ pathway and the PI3K-FOXO signaling pathway [20],[58],[59],[60]. Similarly, crosstalk between other signal transduction pathways has been frequently described, potentially enabling specific ligands for one pathway to modify activity of another signaling pathway [61],[62],[63].

### Defining a pathway activity profile for pluripotency

Activity of signal transduction pathways that are essential for maintaining pluripotency would be expected to decrease upon differentiation. Differentiation of pluripotent hES cells in the mesodermal/endodermal direction was associated with rapid initial decrease (within 1 day) in pathway activity scores for the MAPK-AP1, PI3K, Notch, TGFβ, HH and NFκB pathways, indicating their activity in pluripotent stem cells and supporting a role for a coordinated activity of these signaling pathways in pluripotency maintenance. Of note, ectodermal differentiation data were only available after two weeks of iPS cell culture, and therefore not useful for the purpose of defining pluripotency pathways.

In line with our results, activity of the PI3K, MAPK, Notch, TGFβ, and NFκB pathways has been shown to be necessary for the maintenance of pluripotency [19],[69],[70],[71],[72],[73],[74]. With respect to the HH pathway, all pluripotent cell lines had high pathway activation scores. The HH pathway is known to be important for embryonic development [70]. The expression of key HH signaling molecules in pluripotent stem cells has been reported, but information on functional pathway activity is lacking [71]. HH pathway activity is determined by the interaction between activating and inhibitory GLI transcription factors and this makes it difficult to infer activity of this pathway from the expression of its signaling cascade proteins and transcription factors [31]. Given the currents results it seems plausible that activity of this embryonic pathway is essential to pluripotency. Low Wnt pathway activity scores are fully in line with current thinking on the role of Wnt in human pluripotent stem cells [57]. While we cannot exclude that other signaling pathways may contribute to stem cell pluripotency, based on the current results, we tentatively propose to define a stem cell pluripotency profile as active MAPK-AP1, PI3K, Notch, TGFβ, HH and NFκB pathways; low score for Wnt pathway. We hypothesize that measuring combined activity of these pathways using STP analysis can be used to define and quantify the pluripotency state of a stem cell culture. While Affymetrix-based pluripotency assessment is not very suitable for implementation in day-to-day research, the recent conversion of Affymetrix STP analysis to a platform of qPCR-based STP assays enables generation of a full STP activity or pluripotency profile of a stem cell sample within a few hours on standard lab equipment [72]. This is also expected to facilitate experimental modification of the STP activity profile by controlled manipulation of culture conditions, for example by adding specific pathway ligands, and subsequently analyze the relation between different STP activity profiles and pluripotent stem cell differentiation capability.

### Differentiation-induced changes in signaling pathway activity

The available datasets enabled investigation of changes in STP activity associated with hES cell differentiation to mesoderm, definitive endoderm, and digestive tract precursor cells, and neural progenitor cells, while hES-derived intestinal precursor cells could be compared with adult intestinal crypt stem cells.

Differentiation of hES cells resulted in pathway activity profiles characteristic for the two germ layers [40]. Low Wnt pathway activity in definitive endoderm versus higher Wnt pathway activity scores in mesodermal differentiation is in agreement with a role for the Wnt pathway in mesodermal differentiation and differentiation to mesoderm-derived cardiomyocytes [12],[57],[73],[79],[80]. The simultaneous increase in TGFβ pathway activity in mesodermal differentiation is in line with the required BMP signaling [74],[76]. The return to high HH pathway activity scores upon differentiation to anterior/posterior/mid/hind-gut is in line with the broad role of this pathway in embryonic development of the digestive tract [77].

Specific differentiation to mid/hindgut, as precursor for intestinal cell development, was associated with a marked increase in Wnt and Notch pathway activity and low MAPK pathway activity scores. A comparable STP activity profile (although even higher Wnt pathway activity) was found in adult intestinal crypt stem cells and disappeared upon differentiation to various mucosa cell types. These results underscore the important role of these pathways in early development as well as in adult intestinal physiology [56],[83],[84],[80]. The increase in ER pathway activity scores seen during mucosal differentiation is in line with the role of ER in mucosal physiology [81]. Finally, differentiation of iPS cells to neuronal cells was associated with gain of activity of the Notch pathway, in agreement with its role as a neuronal progenitor signaling pathway [87]. Taken together, these results provide evidence that STP analysis can be used to assess stem cell differentiation state, probably even in a quantitative manner.

### Summary and perspective

The STP assay platform allows quantification of activity of multiple signaling pathways simultaneously in a cell sample. The STP pluripotency profile can be used to phenotypically compare genetically different pluripotent stem cell lines and investigate the relation with differentiation capability. The STP activity profile provides actionable information that can be used to restore or modify pluripotency or cultured cells. The recent conversion to qPCR-based STP analysis facilitates implementation in a research laboratory. The use of STP activity analysis is expected to increase experimental reproducibility across laboratories, and improve standardization of the stem cell field, as well as providing an additional method for quality control in the production of cells for regenerative medicine purposes.

## Author contributions

Y.W.-R.: data analysis, figures, editing

L.H.: development MAPK pathway model, data analysis, figures, editing

W.V.: pathway models, figures, editing

A.v.d.S: concept, data analysis, writing

## Acknowledgements

For the use of dataset GSE17312 we acknowledge the NIH Roadmap Epigenomics Mapping Consortium and their data suppliers, http://nihroadmap.nih.gov/epigenomics/. For the other public datasets analyzed in this publication, we wish to thank all suppliers of these data for their contribution to the open access GEO database.

## SUPPLEMENTARY MATERIALS AND METHODS

### Development of an assay to measure MAPK-AP1 pathway activity

The method to develop assays to measure the functional activity of signal transduction pathways has been described in detail before [16]. The mathematical approach to develop Bayesian network models for the measurement of signal transduction pathway activity has been described [16]. In brief, a Bayesian computational network model for a signal transduction pathway is created to infer the probability that the pathway-associated transcription factor is in a transcriptionally active state. The Bayesian network describes (i) the causal relation that a target gene is up- or downregulated depending on the transcription complex being active or inactive and (ii) the causal relation that a signal on an Affymetrix HG-U133 Plus 2.0 probeset is high or low depending on the target gene being up or down. These relations are probabilistic; the parameters describing relation (i) are based on literature evidence, and the parameters describing relation (ii) are based on calibration data of samples with ground truth information about their pathway activity state.

*Supplementary Table S1. List with MAPK-AP1 target genes according to HGNC approved symbols used for development of the pathway activity assay.*

BCL2L11, CCND1, DDIT3, DNMT1, EGFR, ENPP2, EZR, FASLG, VEGFD, GLRX, IL2, IVL, LORICRIN, MMP1, MMP3, MMP9, PLAU, PLAUR, PTGS2, SERPINE1, SNCG, TIMP1, TP53, VIM.

---

**Supplementary Figure S1.**
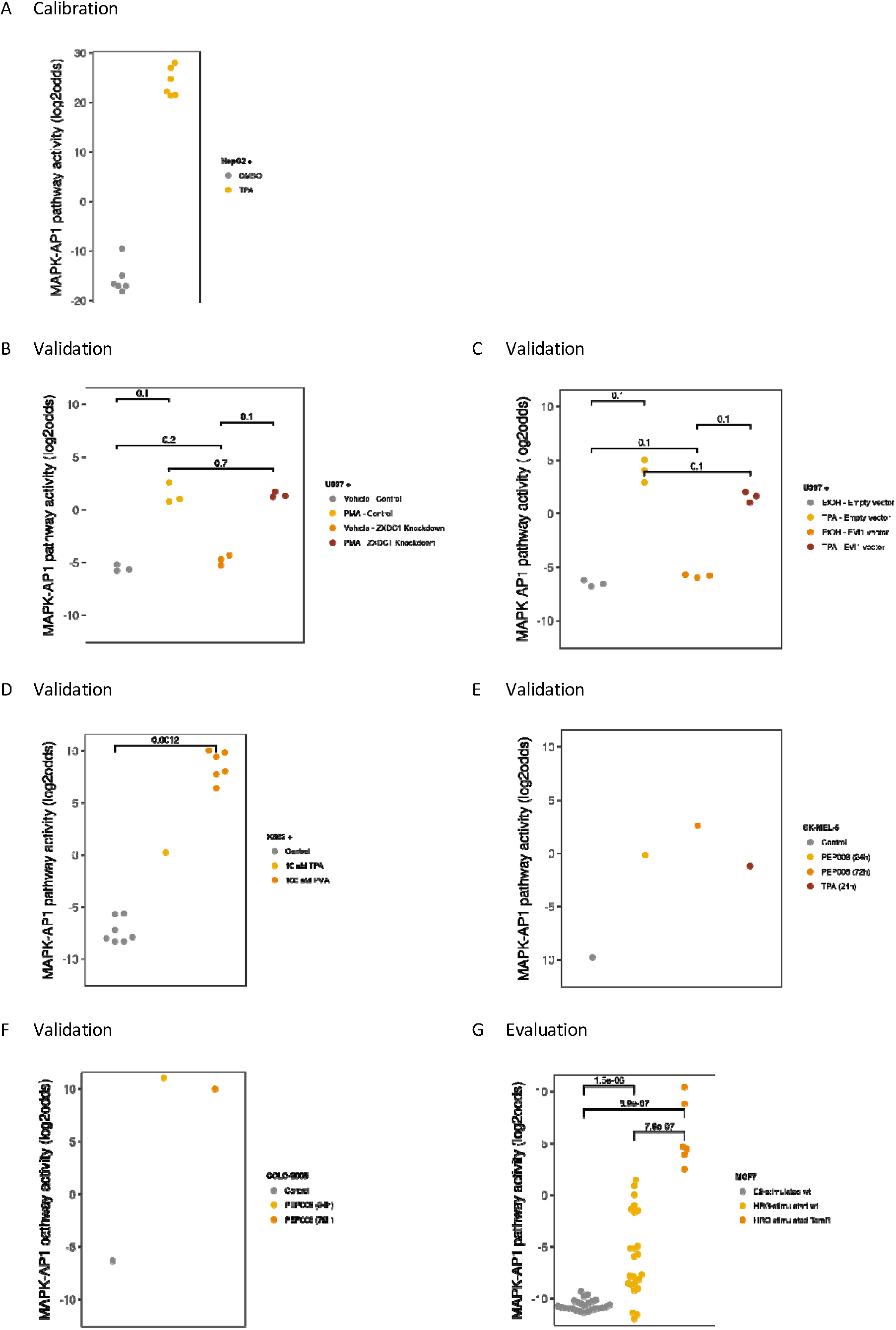
Calibration and validation of the MAPK-AP1 pathway assay. For calibration purposes vehicle-treated (MAPK-AP1 inactive) and 500 nM TPA-treated (MAPK-AP1 active) HepG2 liver cell samples were used (A) from GSE28878 [49]. After freezing the model, independent samples were used for validation (B-F) and evaluation of the model (G). B. GSE45417, 100 nM TPA-treated U937 cells [50]; C. GSE66853, U937 cells were treated with TPA in the absence or presence of an EVI1 expression vector to activate the MAPK-AP1 signaling pathway [51]; D. E-MEXP-2213 and E-MEXP-2573 (ArrayExpress database; https://www.ebi.ac.uk/arrayexpress/), K562 erythroleukemia cells were treated with 10 and 100 nM TPA/PMA, respectively [52], [53]; E. GSE8742, SK-MEL-5 (melanoma) cells and F. GSE8742, COLO-2005 (colon cancer; Supplementary Figure S1F) cells were treated with 1 μg/mL PEP008 or 1000 ng/mL TPA (only the SK-MEL-5 cells) or vehicle for 24h, followed by 72h recovery [54]; G. GSE21618, wild-type (wt) MCF7 human breast cancer cells and tamoxifen-resistant MCF7 were treated with 100 nM 17beta-estradiol (E2) or 10 nM growth hormone heregulin (HRG) to activate the MAPK-AP1 signaling pathway [55].

**Supplementary Figure S2.**
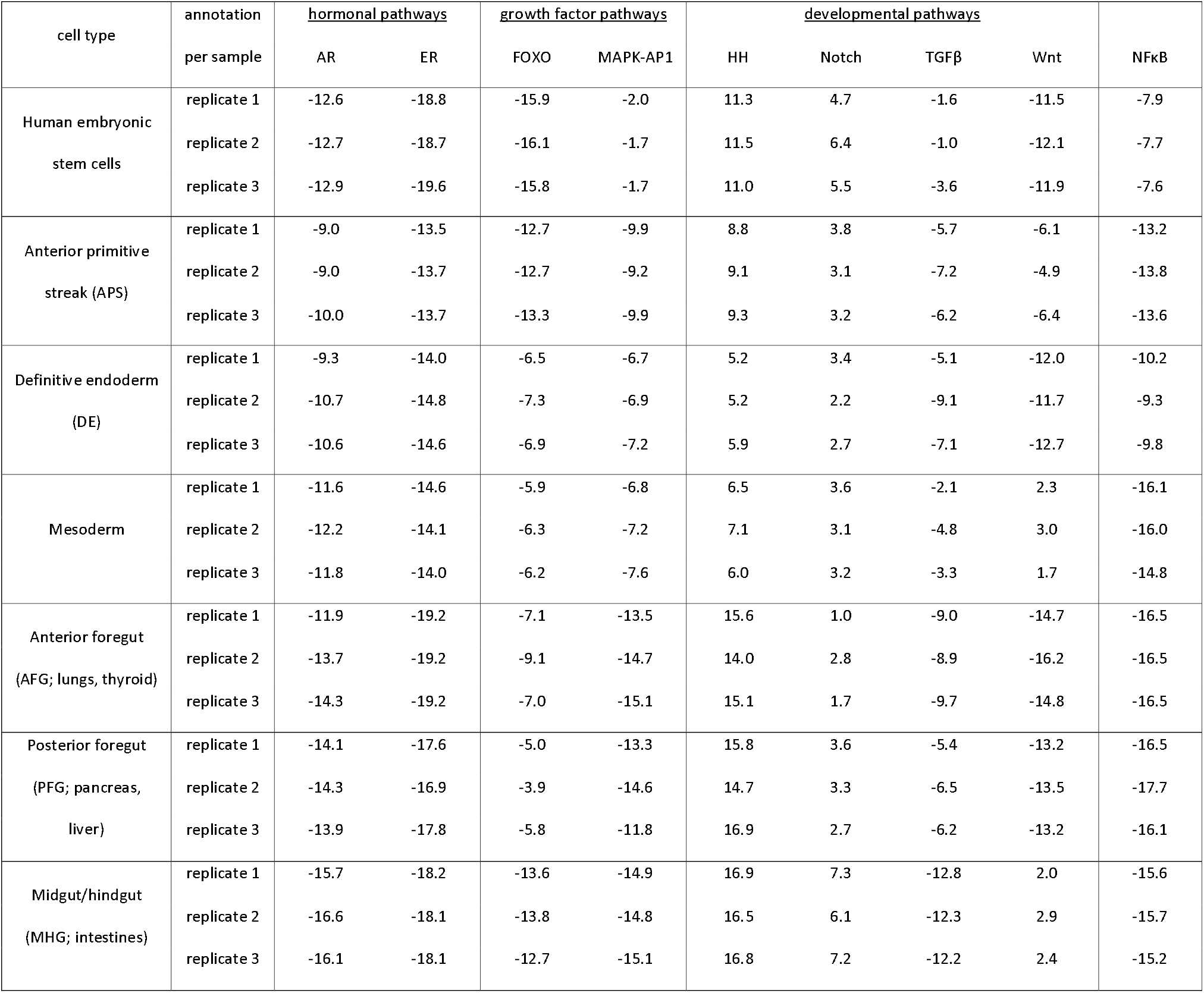
Full signaling pathway analysis data per sample, corresponding to the results in Figure 2 (dataset GSE52658 [40]).

**Supplementary Figure S3.**
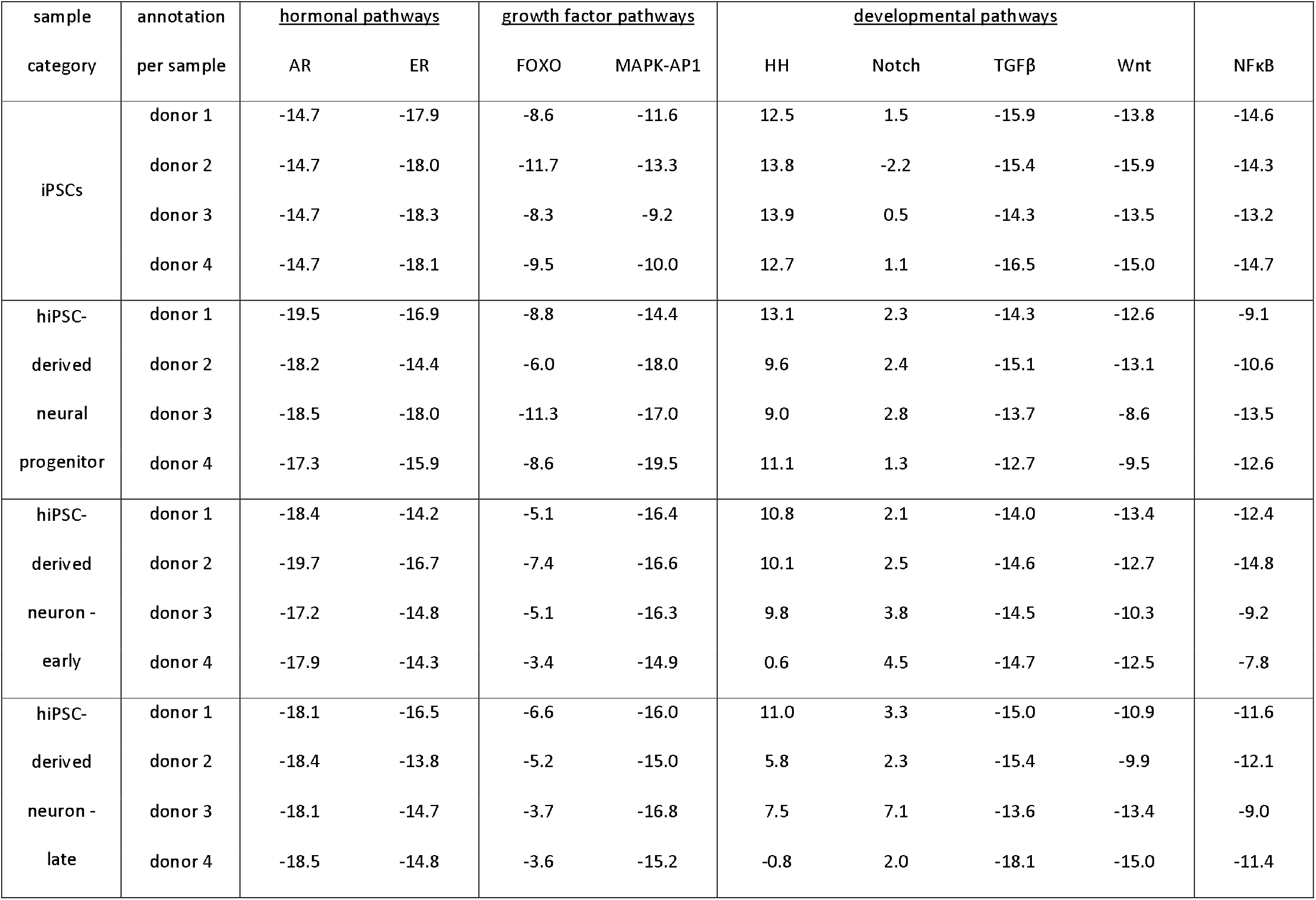
Full signaling pathway analysis data per sample, corresponding to the results presented in Figure 3 (dataset GSE74358 [45]).

## References

[1] C. L. Mummery, A. van de Stolpe, B. Roelen, and H. Clevers, Stem Cells: Scientific Facts and Fiction. Waltham, 2021.

[2] “Human Stem Cell Manual - 2nd Edition.” https://www.elsevier.com/books/human-stem-cell-manual/peterson/978-0-12-385473-5 (accessed Apr. 13, 2021).

[3] C. Mummery, A. Van de Stolpe, B. A. J. Roelen, and H. Clevers, Stem Cells: Scientific Facts and Fiction, 2nd ed. Academic Press, 2014.

[4] S. Dimmeler, S. Ding, T. A. Rando, and A. Trounson, “Translational strategies and challenges in regenerative medicine,” Nat. Med., vol. 20, no. 8, pp. 814–821, Aug. 2014, doi: 10.1038/nm.3627.

[5] E. A. Kimbrel and R. Lanza, “Current status of pluripotent stem cells: moving the first therapies to the clinic,” Nat. Rev. Drug Discov., vol. 14, no. 10, pp. 681–692, Oct. 2015, doi: 10.1038/nrd4738.

[6] S. J. Forbes and N. Rosenthal, “Preparing the ground for tissue regeneration: from mechanism to therapy,” Nat. Med., vol. 20, no. 8, pp. 857–869, Aug. 2014, doi: 10.1038/nm.3653.

[7] A. van de Stolpe and J. den Toonder, “Workshop meeting report Organs-on-Chips: human disease models,” Lab. Chip, vol. 13, no. 18, pp. 3449–3470, Sep. 2013, doi: 10.1039/c3lc50248a.

[8] S. Badylak and N. Rosenthal, “Regenerative medicine: are we there yet?,” Npj Regen. Med., vol. 2, no. 1, p. 2, Jan. 2017, doi: 10.1038/s41536-016-0005-9.

[9] S. Dakhore, B. Nayer, and K. Hasegawa, “Human Pluripotent Stem Cell Culture: Current Status, Challenges, and Advancement,” Stem Cells Int., vol. 2018, Nov. 2018, doi: 10.1155/2018/7396905.

[10] V. Schwach et al., “A COUP-TFII Human Embryonic Stem Cell Reporter Line to Identify and Select Atrial Cardiomyocytes,” Stem Cell Rep., vol. 9, no. 6, pp. 1765–1779, Dec. 2017, doi: 10.1016/j.stemcr.2017.10.024.

[11] X. Bao et al., “Gene Editing to Generate Versatile Human Pluripotent Stem Cell Reporter Lines for Analysis of Differentiation and Lineage Tracing,” Stem Cells Dayt. Ohio, vol. 37, no. 12, pp. 1556–1566, Dec. 2019, doi: 10.1002/stem.3096.

[12] S. C. Den Hartogh et al., “Dual reporter MESP1 mCherry/w-NKX2-5 eGFP/w hESCs enable studying early human cardiac differentiation,” Stem Cells Dayt. Ohio, vol. 33, no. 1, pp. 56–67, Jan. 2015, doi: 10.1002/stem.1842.

[13] S. Sullivan et al., “Quality control guidelines for clinical-grade human induced pluripotent stem cell lines,” Regen. Med., vol. 13, no. 7, pp. 859–866, 2018, doi: 10.2217/rme-2018-0095.

[14] D. Ortmann and L. Vallier, “Variability of human pluripotent stem cell lines,” Curr. Opin. Genet. Dev., vol. 46, pp. 179–185, Oct. 2017, doi: 10.1016/j.gde.2017.07.004.

[15] D. Zheng, X. Wang, and R.-H. Xu, “Concise Review: One Stone for Multiple Birds: Generating Universally Compatible Human Embryonic Stem Cells,” Stem Cells Dayt. Ohio, vol. 34, no. 9, pp. 2269–2275, 2016, doi: 10.1002/stem.2407.

[16] W. Verhaegh et al., “Selection of personalized patient therapy through the use of knowledge-based computational models that identify tumor-driving signal transduction pathways,” Cancer Res., vol. 74, no. 11, pp. 2936–2945, Jun. 2014, doi: 10.1158/0008-5472.CAN-13-2515.

[17] C. A. Chacón-Martínez, J. Koester, and S. A. Wickström, “Signaling in the stem cell niche: regulating cell fate, function and plasticity,” Development, vol. 145, no. 15, p. dev165399, Aug. 2018, doi: 10.1242/dev.165399.

[18] L. Cantley, T. Hunter, R. Sever, and J. Thorner, Signal Transduction: Principles, Pathways, and Processes. New York: Cold Spring Harbor Laboratory Press, 2014.

[19] J. S. L. Yu and W. Cui, “Proliferation, survival and metabolism: the role of PI3K/AKT/mTOR signalling in pluripotency and cell fate determination,” Development, vol. 143, no. 17, pp. 3050–3060, Sep. 2016, doi: 10.1242/dev.137075.

[20] F. Haghighi et al., “bFGF-mediated pluripotency maintenance in human induced pluripotent stem cells is associated with NRAS-MAPK signaling,” Cell Commun. Signal. CCS, vol. 16, no. 1, p. 96, 05 2018, doi: 10.1186/s12964-018-0307-1.

[21] J.-F. Arnal et al., “Membrane and Nuclear Estrogen Receptor Alpha Actions: From Tissue Specificity to Medical Implications,” Physiol. Rev., vol. 97, no. 3, pp. 1045–1087, May 2017, doi: 10.1152/physrev.00024.2016.

[22] M. Kono, T. Fujii, B. Lim, M. S. Karuturi, D. Tripathy, and N. T. Ueno, “Androgen Receptor Function and Androgen Receptor-Targeted Therapies in Breast Cancer: A Review,” JAMA Oncol., vol. 3, no. 9, pp. 1266–1273, Sep. 2017, doi: 10.1001/jamaoncol.2016.4975.

[23] C. Siebel and U. Lendahl, “Notch Signaling in Development, Tissue Homeostasis, and Disease,” Physiol. Rev., vol. 97, no. 4, pp. 1235–1294, Aug. 2017, doi: 10.1152/physrev.00005.2017.

[24] M. Burotto, V. L. Chiou, J.-M. Lee, and E. C. Kohn, “The MAPK pathway across different malignancies: a new perspective,” Cancer, vol. 120, no. 22, pp. 3446–3456, Nov. 2014, doi: 10.1002/cncr.28864.

[25] E. Pardali, G. Sanchez-Duffhues, M. C. Gomez-Puerto, and P. Ten Dijke, “TGF-β-Induced Endothelial-Mesenchymal Transition in Fibrotic Diseases,” Int. J. Mol. Sci., vol. 18, no. 10, Oct. 2017, doi: 10.3390/ijms18102157.

[26] G. Sánchez-Duffhues, A. G. de Vinuesa, and P. ten Dijke, “Endothelial-to-mesenchymal transition in cardiovascular diseases: Developmental signaling pathways gone awry,” Dev. Dyn., vol. 247, no. 3, pp. 492–508, 2018, doi: 10.1002/dvdy.24589.

[27] F. A. High and J. A. Epstein, “The multifaceted role of Notch in cardiac development and disease,” Nat. Rev. Genet., vol. 9, no. 1, pp. 49–61, Jan. 2008, doi: 10.1038/nrg2279.

[28] G. Sanchez-Duffhues, V. Orlova, and P. Ten Dijke, “In Brief: Endothelial-to-mesenchymal transition,” J. Pathol., vol. 238, no. 3, pp. 378–380, Feb. 2016, doi: 10.1002/path.4653.

[29] M. Bienz and H. Clevers, “Linking colorectal cancer to Wnt signaling,” Cell, vol. 103, no. 2, pp. 311–320, Oct. 2000, doi: 10.1016/s0092-8674(00)00122-7.

[30] S. S. Karhadkar et al., “Hedgehog signalling in prostate regeneration, neoplasia and metastasis,” Nature, vol. 431, no. 7009, pp. 707–712, Oct. 2004, doi: 10.1038/nature02962.

[31] J. Taipale and P. A. Beachy, “The Hedgehog and Wnt signalling pathways in cancer,” Nature, vol. 411, no. 6835, pp. 349–354, May 2001, doi: 10.1038/35077219.

[32] D. Hanahan and R. A. Weinberg, “Hallmarks of cancer: the next generation,” Cell, vol. 144, no. 5, pp. 646–674, Mar. 2011, doi: 10.1016/j.cell.2011.02.013.

[33] A. van de Stolpe, “On the origin and destination of cancer stem cells: a conceptual evaluation,” Am. J. Cancer Res., vol. 3, no. 1, pp. 107–116, Jan. 2013.

[34] W. Verhaegh and A. Van de Stolpe, “Knowledge-based computational models,” Oncotarget, vol. 5, no. 14, pp. 5196–5197, Jul. 2014, doi: 10.18632/oncotarget.2276.

[35] H. van Ooijen et al., “Assessment of Functional Phosphatidylinositol 3-Kinase Pathway Activity in Cancer Tissue Using Forkhead Box-O Target Gene Expression in a Knowledge-Based Computational Model,” Am. J. Pathol., vol. 188, no. 9, pp. 1956–1972, Sep. 2018, doi: 10.1016/j.ajpath.2018.05.020.

[36] K. Canté-Barrett et al., “A Molecular Test for Quantifying Functional Notch Signaling Pathway Activity in Human Cancer,” Cancers, vol. 12, no. 11, Art. no. 11, Nov. 2020, doi: 10.3390/cancers12113142.

[37] A. van de Stolpe, L. Holtzer, H. van Ooijen, M. A. de Inda, and W. Verhaegh, “Enabling precision medicine by unravelling disease pathophysiology: quantifying signal transduction pathway activity across cell and tissue types,” Sci. Rep., vol. 9, no. 1, p. 1603, Feb. 2019, doi: 10.1038/s41598-018-38179-x.

[38] A. van de Stolpe, “Quantitative Measurement of Functional Activity of the PI3K Signaling Pathway in Cancer,” Cancers, vol. 11, no. 3, Mar. 2019, doi: 10.3390/cancers11030293.

[39] Z.-Y. Tsai, S. Singh, S.-L. Yu, C.-H. Chou, and S. S.-L. Li, “A feeder-free culture using autogeneic conditioned medium for undifferentiated growth of human embryonic stem cells: comparative expression profiles of mRNAs, microRNAs and proteins among different feeders and conditioned media,” BMC Cell Biol., vol. 11, p. 76, Oct. 2010, doi: 10.1186/1471-2121-11-76.

[40] K. M. Loh et al., “Efficient endoderm induction from human pluripotent stem cells by logically directing signals controlling lineage bifurcations,” Cell Stem Cell, vol. 14, no. 2, pp. 237–252, Feb. 2014, doi: 10.1016/j.stem.2013.12.007.

[41] K. Giraud-Triboult, C. Rochon-Beaucourt, X. Nissan, B. Champon, S. Aubert, and G. Piétu, “Combined mRNA and microRNA profiling reveals that miR-148a and miR-20b control human mesenchymal stem cell phenotype via EPAS1,” Physiol. Genomics, vol. 43, no. 2, pp. 77–86, Jan. 2011, doi: 10.1152/physiolgenomics.00077.2010.

[42] S. S.-L. Li et al., “Target identification of microRNAs expressed highly in human embryonic stem cells,” J. Cell. Biochem., vol. 106, no. 6, pp. 1020–1030, Apr. 2009, doi: 10.1002/jcb.22084.

[43] B. E. Bernstein et al., “The NIH Roadmap Epigenomics Mapping Consortium,” Nat. Biotechnol., vol. 28, no. 10, pp. 1045–1048, Oct. 2010, doi: 10.1038/nbt1010-1045.

[44] V. Amin et al., “Epigenomic footprints across 111 reference epigenomes reveal tissue-specific epigenetic regulation of lincRNAs,” Nat. Commun., vol. 6, p. 6370, Feb. 2015, doi: 10.1038/ncomms7370.

[45] K. H. Kim et al., “Transcriptomic Analysis of Induced Pluripotent Stem Cells Derived from Patients with Bipolar Disorder from an Old Order Amish Pedigree,” PLOS ONE, vol. 10, no. 11, p. e0142693, Nov. 2015, doi: 10.1371/journal.pone.0142693.

[46] E. J. van Zoelen, I. Duarte, J. M. Hendriks, and S. P. van der Woning, “TGFβ-induced switch from adipogenic to osteogenic differentiation of human mesenchymal stem cells: identification of drug targets for prevention of fat cell differentiation,” Stem Cell Res. Ther., vol. 7, no. 1, p. 123, Aug. 2016, doi: 10.1186/s13287-016-0375-3.

[47] P. Jung et al., “Isolation and in vitro expansion of human colonic stem cells,” Nat. Med., vol. 17, no. 10, pp. 1225–1227, Sep. 2011, doi: 10.1038/nm.2470.

[48] M. Karin, “The Regulation of AP-1 Activity by Mitogen-activated Protein Kinases,” J. Biol. Chem., vol. 270, no. 28, pp. 16483–16486, Jul. 1995, doi: 10.1074/jbc.270.28.16483.

[49] C. Magkoufopoulou, S. M. H. Claessen, M. Tsamou, D. G. J. Jennen, J. C. S. Kleinjans, and J. H. M. van Delft, “A transcriptomics-based in vitro assay for predicting chemical genotoxicity in vivo,” Carcinogenesis, vol. 33, no. 7, pp. 1421–1429, Jul. 2012, doi: 10.1093/carcin/bgs182.

[50] J. E. Ramsey and J. D. Fontes, “The zinc finger transcription factor ZXDC activates CCL2 gene expression by opposing BCL6-mediated repression,” Mol. Immunol., vol. 56, no. 4, pp. 768–780, Dec. 2013, doi: 10.1016/j.molimm.2013.07.001.

[51] B. Steinmetz et al., “The oncogene EVI1 enhances transcriptional and biological responses of human myeloid cells to all-trans retinoic acid,” Cell Cycle, vol. 13, no. 18, pp. 2931–2943, Sep. 2014, doi: 10.4161/15384101.2014.946869.

[52] F. Navarro et al., “miR-34a contributes to megakaryocytic differentiation of K562 cells independently of p53,” Blood, vol. 114, no. 10, pp. 2181–2192, Sep. 2009, doi: 10.1182/blood-2009-02-205062.

[53] S. J. Goodfellow et al., “WT1 and its transcriptional cofactor BASP1 redirect the differentiation pathway of an established blood cell line,” Biochem. J., vol. 435, no. 1, pp. 113–125, Apr. 2011, doi: 10.1042/BJ20101734.

[54] S. A. Mason, S.-J. Cozzi, C. J. Pierce, S. J. Pavey, P. G. Parsons, and G. M. Boyle, “The induction of senescence-like growth arrest by protein kinase C-activating diterpene esters in solid tumor cells,” Invest. New Drugs, vol. 28, no. 5, pp. 575–586, Oct. 2010, doi: 10.1007/s10637-009-9292-y.

[55] M. Oyama et al., “Integrated quantitative analysis of the phosphoproteome and transcriptome in tamoxifen-resistant breast cancer,” J. Biol. Chem., vol. 286, no. 1, pp. 818–829, Jan. 2011, doi: 10.1074/jbc.M110.156877.

[56] H. Clevers, “The intestinal crypt, a prototype stem cell compartment,” Cell, vol. 154, no. 2, pp. 274–284, Jul. 2013, doi: 10.1016/j.cell.2013.07.004.

[57] A. de Jaime-Soguero, W. A. Abreu de Oliveira, and F. Lluis, “The Pleiotropic Effects of the Canonical Wnt Pathway in Early Development and Pluripotency,” Genes, vol. 9, no. 2, Feb. 2018, doi: 10.3390/genes9020093.

[58] A. Sundqvist et al., “JUNB governs a feed-forward network of TGFβ signaling that aggravates breast cancer invasion,” Nucleic Acids Res., vol. 46, no. 3, pp. 1180–1195, Feb. 2018, doi: 10.1093/nar/gkx1190.

[59] E. Aksamitiene, A. Kiyatkin, and B. N. Kholodenko, “Cross-talk between mitogenic Ras/MAPK and survival PI3K/Akt pathways: a fine balance,” Biochem. Soc. Trans., vol. 40, no. 1, pp. 139–146, Feb. 2012, doi: 10.1042/BST20110609.

[60] M. C. Mendoza, E. E. Er, and J. Blenis, “The Ras-ERK and PI3K-mTOR pathways: cross-talk and compensation,” Trends Biochem. Sci., vol. 36, no. 6, pp. 320–328, Jun. 2011, doi: 10.1016/j.tibs.2011.03.006.

[61] Y. Katoh and M. Katoh, “Hedgehog target genes: mechanisms of carcinogenesis induced by aberrant hedgehog signaling activation,” Curr. Mol. Med., vol. 9, no. 7, pp. 873–886, Sep. 2009.

[62] H. Nishi, E. Demir, and A. R. Panchenko, “Crosstalk between signaling pathways provided by single and multiple protein phosphorylation sites,” J. Mol. Biol., vol. 427, no. 2, pp. 511–520, Jan. 2015, doi: 10.1016/j.jmb.2014.11.001.

[63] K. Luo, “Signaling Cross Talk between TGF-β/Smad and Other Signaling Pathways,” Cold Spring Harb. Perspect. Biol., vol. 9, no. 1, Jan. 2017, doi: 10.1101/cshperspect.a022137.

[64] A. C. Mullen and J. L. Wrana, “TGF-β Family Signaling in Embryonic and Somatic Stem-Cell Renewal and Differentiation,” Cold Spring Harb. Perspect. Biol., vol. 9, no. 7, Jul. 2017, doi: 10.1101/cshperspect.a022186.

[65] O. Takase et al., “The Role of NF-κB Signaling in the Maintenance of Pluripotency of Human Induced Pluripotent Stem Cells,” PLoS ONE, vol. 8, no. 2, Feb. 2013, doi: 10.1371/journal.pone.0056399.

[66] X. Yu et al., “Notch signaling activation in human embryonic stem cells is required for embryonic but not trophoblastic lineage commitment,” Cell Stem Cell, vol. 2, no. 5, pp. 461–471, May 2008, doi: 10.1016/j.stem.2008.03.001.

[67] U. Koch, R. Lehal, and F. Radtke, “Stem cells living with a Notch,” Development, vol. 140, no. 4, pp. 689–704, Feb. 2013, doi: 10.1242/dev.080614.

[68] W. Li et al., “Rapid induction and long-term self-renewal of primitive neural precursors from human embryonic stem cells by small molecule inhibitors,” Proc. Natl. Acad. Sci. U. S. A., vol. 108, no. 20, pp. 8299–8304, May 2011, doi: 10.1073/pnas.1014041108.

[69] E. Ben-Shushan, E. Feldman, and B. E. Reubinoff, “Notch signaling regulates motor neuron differentiation of human embryonic stem cells,” Stem Cells Dayt. Ohio, vol. 33, no. 2, pp. 403–415, Feb. 2015, doi: 10.1002/stem.1873.

[70] A. P. McMahon, P. W. Ingham, and C. J. Tabin, “Developmental roles and clinical significance of hedgehog signaling,” Curr. Top. Dev. Biol., vol. 53, pp. 1–114, 2003, doi: 10.1016/s0070-2153(03)53002-2.

[71] S. M. Wu, A. B. H. Choo, M. G. S. Yap, and K. K.-K. Chan, “Role of Sonic hedgehog signaling and the expression of its components in human embryonic stem cells,” Stem Cell Res., vol. 4, no. 1, pp. 38–49, Jan. 2010, doi: 10.1016/j.scr.2009.09.002.

[72] A. van de Stolpe et al., “RNA Based Approaches to Profile Oncogenic Pathways From Low Quantity Samples to Drive Precision Oncology Strategies,” Front. Genet., vol. 11, 2021, doi: 10.3389/fgene.2020.598118.

[73] J. Y. Tan, G. Sriram, A. J. Rufaihah, K. G. Neoh, and T. Cao, “Efficient derivation of lateral plate and paraxial mesoderm subtypes from human embryonic stem cells through GSKi-mediated differentiation,” Stem Cells Dev., vol. 22, no. 13, pp. 1893–1906, Jul. 2013, doi: 10.1089/scd.2012.0590.

[74] J. Rao et al., “Stepwise Clearance of Repressive Roadblocks Drives Cardiac Induction in Human ESCs,” Cell Stem Cell, vol. 18, no. 3, pp. 341–353, Mar. 2016, doi: 10.1016/j.stem.2015.11.019.

[75] X. Lian et al., “Robust cardiomyocyte differentiation from human pluripotent stem cells via temporal modulation of canonical Wnt signaling,” Proc. Natl. Acad. Sci. U. S. A., vol. 109, no. 27, pp. E1848–1857, Jul. 2012, doi: 10.1073/pnas.1200250109.

[76] V. V. Orlova, S. Chuva de Sousa Lopes, and G. Valdimarsdottir, “BMP-SMAD signaling: From pluripotent stem cells to cardiovascular commitment,” Cytokine Growth Factor Rev., vol. 27, pp. 55–63, Feb. 2016, doi: 10.1016/j.cytogfr.2015.11.007.

[77] K. Fukuda and S. Yasugi, “Versatile roles for sonic hedgehog in gut development,” J. Gastroenterol., vol. 37, no. 4, pp. 239–246, 2002, doi: 10.1007/s005350200030.

[78] G. M. Collu, A. Hidalgo-Sastre, and K. Brennan, “Wnt-Notch signalling crosstalk in development and disease,” Cell. Mol. Life Sci. CMLS, vol. 71, no. 18, pp. 3553–3567, Sep. 2014, doi: 10.1007/s00018-014-1644-x.

[79] J. R. Spence et al., “Directed differentiation of human pluripotent stem cells into intestinal tissue in vitro,” Nature, vol. 470, no. 7332, pp. 105–109, Feb. 2011, doi: 10.1038/nature09691.

[80] G. Wei et al., “Erk and MAPK signaling is essential for intestinal development through Wnt pathway modulation,” Dev. Camb. Engl., vol. 147, no. 17, Sep. 2020, doi: 10.1242/dev.185678.

[81] C. Zhao, K. Dahlman-Wright, and J.-Å. Gustafsson, “Estrogen Signaling via Estrogen Receptor β,” J. Biol. Chem., vol. 285, no. 51, pp. 39575–39579, Dec. 2010, doi: 10.1074/jbc.R110.180109.

[82] L. Grandbarbe, J. Bouissac, M. Rand, M. H. de Angelis, S. Artavanis-Tsakonas, and E. Mohier, “Delta-Notch signaling controls the generation of neurons/glia from neural stem cells in a stepwise process,” Development, vol. 130, no. 7, pp. 1391–1402, Apr. 2003, doi: 10.1242/dev.00374.

[83] Y. Hirabayashi et al., “The Wnt/β-catenin pathway directs neuronal differentiation of cortical neural precursor cells,” Development, vol. 131, no. 12, pp. 2791–2801, Jun. 2004, doi: 10.1242/dev.01165.

[84] A. van de Stolpe, L. Holtzer, H. van Ooijen, M. A. de Inda, and W. Verhaegh, “Enabling precision medicine by unravelling disease pathophysiology: quantifying signal transduction pathway activity across cell and tissue types,” Sci. Rep., vol. 9, no. 1, p. 1603, Feb. 2019, doi: 10.1038/s41598-018-38179-x.

